# EMDA: A Python package for Electron Microscopy Data Analysis

**DOI:** 10.1101/2021.07.26.453750

**Authors:** Rangana Warshamanage, Keitaro Yamashita, Garib N. Murshudov

## Abstract

An open-source Python library EMDA for cryo-EM map and model manipulation is presented with a specific focus on validation. The use of several functionalities in the library is presented through several examples. The utility of local correlation as a metric for identifying map-model differences and unmodeled regions in maps, and how it is used as a metric of map-model validation is demonstrated. The mapping of local correlation to individual atoms, and its use to draw insights on local signal variations are discussed. EMDA’s likelihood-based map overlay is demonstrated by carrying out a superposition of two domains in two related structures. The overlay is carried out first to bring both maps into the same coordinate frame and then to estimate the relative movement of domains. Finally, the map magnification refinement in EMDA is presented with an example to highlight the importance of adjusting the map magnification in structural comparison studies.

## 1. Introduction

Single-particle cryo-electron microscopy (cryo-EM) has become an increasingly popular structure determination tool among structural biologists (Faruqi and McMullan, 2011; Kühlbrandt, 2014; Lyumkis, 2019; Lyumkis et al., 2013; Scheres, 2014). The technique has evolved at an unprecedented speed in the past few years as shown by the rapid growth of cryo-EM structure depositions into the Electron Microscopy Data Bank – EMDB (Lawson et al., 2020). As the number of depositions into EMDB increases, it is important to maintain quality standards for both maps and atomic models not only to ensure their reliability, but also to prevent accumulation of errors.

The EM validation task force 2010 (Henderson et al., 2012) has recognized the critical need of validation standards to assess the quality of EM maps, models and their fits. The task force’s recommendations for map validation included the tilt-pair experiments for the absolute hand determination (Rosenthal and Henderson, 2003), the raw image to 3D structure projection matching for validating reconstruction accuracy and the data coverage (Orlova et al., 1996; Tang et al., 2007), statistical tests using map variances for assessing the map quality and interpretability (Ménétret et al., 2007; Penczek et al., 2006), resolution estimation through Fourier Shell Correlation (FSC) using fully independent half data sets (Scheres and Chen, 2012), visual assessment of map features to the claimed resolution, and the identification and validation of the map symmetry where applicable (Reboul et al., 2020).

The task force has identified the model validation in cryo-EM as an area for further research, mainly due to the fact that, at the time, there were few high resolution cryo-EM structures. Thus, the recommendations for model validation included, among others, the assessment of subunits and their interfaces according to the guidelines proposed by the PDB (Read et al., 2011), the assessment of agreement between the model and the map utilizing chemical measures such as chemical properties and atomic interactions and their clashes as employed in EMFIT program (Rossmann, 2000; Rossmann et al., 2001) or statistical measures such as correlation coefficient.

Since the first meeting in 2010, the field has grown by accumulating many methods and tools to address the issue of map validation. Examples include the gold standard FSC to monitor the map overfitting into noise during reconstruction (Rosenthal and Henderson, 2003; Scheres and Chen, 2012), tilt-pair validation to assess the accuracy of initial angle assignment (Wasilewski and Rosenthal, 2014) and the false discovery maps for visual assessment of map features (Beckers et al., 2019).

The progress in atomic model building, refinement and validation has also been substantial. Resolution in cryo-EM reconstructions vary widely, however the progress made in the field of atomic model building encapsulates modelling tools for low, medium to high resolution. Examples include Chimera (Pettersen et al., 2004), DockEM (Roseman, 2000), FlexEM (Topf et al., 2008), COOT(Brown et al., 2015), DireX (Wang and Schröder, 2012), MDFF (Trabuco et al., 2009), Cryo-Fit (Kim et al., 2019), Rosetta (Wang et al., 2016), MDeNM-EMfit (Costa et al., 2020). The atomic model refinement has also gained a significant progress. Unlike in crystallography, cryo-EM maps contain both amplitudes and phases and the atomic model refinement programs can use a phased likelihood target function as employed in REFMAC5 (Murshudov, 2016; Nicholls et al., 2018) or real space target functions as used in phenix.real_space_refine (Afonine et al., 2018a), ISOLDE (Croll, 2018) and COOT (Brown et al., 2015). There has also been a considerable progress in map and model validation. Examples include various metrics for map-model fit validation (Afonine et al., 2018b; Barad et al., 2015; Brown et al., 2015; Pintilie et al., 2020; Ramírez-Aportela et al., 2021), and chemistry and geometry based tools for model validation (Emsley et al., 2010; Prisant et al., 2020). The EM practitioners can access these methods and tools as parts of stand-alone packages, separate tools in collaborative projects such as CCP-EM (Burnley et al., 2017), or as web based tools such as EMDB validation server (https://www.ebi.ac.uk/pdbe/emdb/validation/fsc/), Molprobity server (http://molprobity.biochem.duke.edu/) etc.

Developing better validation methods and tools in cryo-EM is an active area of research because the goal of validation in cryo-EM is a changing target (Lawson et al., 2020). The metrics for validation should evolve as the field progresses towards the atomic resolution because the methods that are applicable to low and medium resolution may not be equally applicable to atomic resolution data and derived models.

In this paper we present Electron Microscopy Data Analytical toolkit (EMDA) - a new Python package for post reconstruction/atomic model refinement analysis and validation of cryo-EM maps and models. EMDA is a portable Python package with a command line and an Application Programming Interface (API) for Python programmers.

While EMDA contains several useful functionalities for cryo-EM map and model manipulation, we describe only three of them in detail in this paper. The local correlation in real space is a metric for detecting the map signal and evaluating the degree of local agreement between an atomic model and a cryo-EM map. We describe the mathematics relevant to correlation calculation in section 3.1. Section 4.1 includes examples demonstrating various uses of local correlation. Map superposition is an important operation in cryo-EM. In structure comparison studies, it brings all maps into a common coordinate frame for comparison. In the difference and average map calculations, the superposition is an essential first step to align the input maps. The available tools for superposing maps include Chimera’s Fit-in-map (Pettersen et al., 2004), TEMPy2 (Cragnolini et al., 2021). EMDA map overlay is based on the maximisation of the likelihood function described in section 3.2. An example demonstrating the overlay is given in section 4.2.

Magnification of an EM map is related to the microscope optics on which the data has been collected. During merging of several data sets collected on different microscopes or on the same microscopy with different optical alignments, their magnifications may need to be adjusted to a reference (Wilkinson et al., 2019). The reference can be another map whose accurate pixel size is already known or an atomic model derived independently of the map whose magnification is sought. The use of EMDA for map magnification correction is demonstrated in section 4.3.

The rest of the paper is organised into three sections. The first section covers the design and the infrastructure of EMDA with an introduction to the EMDA command line and Application Programming Interface (API). The second section describes the mathematical framework of EMDA behind those three functionalities. In the third section, we demonstrate each functionality by examples. Lastly, the conclusions and outlook followed by information about the package availability are given.

## 2. EMDA architecture

EMDA is written primarily in Python using Numpy (Harris et al., 2020), Scipy (Virtanen et al., 2020) and Matplotlib (Hunter, 2007), however, numerically intensive tasks are written in Fortran. F2PY (Peterson, 2009) mediates the communication between Python and Fortran. This combination allows us to integrate powerful numerical calculations with abstraction features in Python.

EMDA code is organized into three layers as shown in Fig. 1. The innermost layer (Layer 3) consists of core and extension modules. The core modules provide basic services such as read & write, format conversion, resampling, binning etc. All higher-level functionalities such as rigid-body fitting, magnification refinement, difference map calculations are provided through extensions. The extension modules use the basic services provided by the core modules. Both core and extension modules are wrapped into another module to form the EMDA API (Layer 2). API abstracts the underlying complexity of the code into methods and objects providing a simplified mechanism for other developers to gain advantage of EMDA infrastructure. EMDA-API functions are further wrapped to form the EMDA command-line-interface (Layer 1). The users can access the underlying functionalities through the command line. Each functionality is callable with a keyword followed by a set of arguments. A list of up-to-date functionalities with their arguments are given in https://emda.readthedocs.io. In addition, a tutorial describing the presented examples in this paper can be found in https://www2.mrc-lmb.cam.ac.uk/groups/murshudov/.

**Fig. 1.**
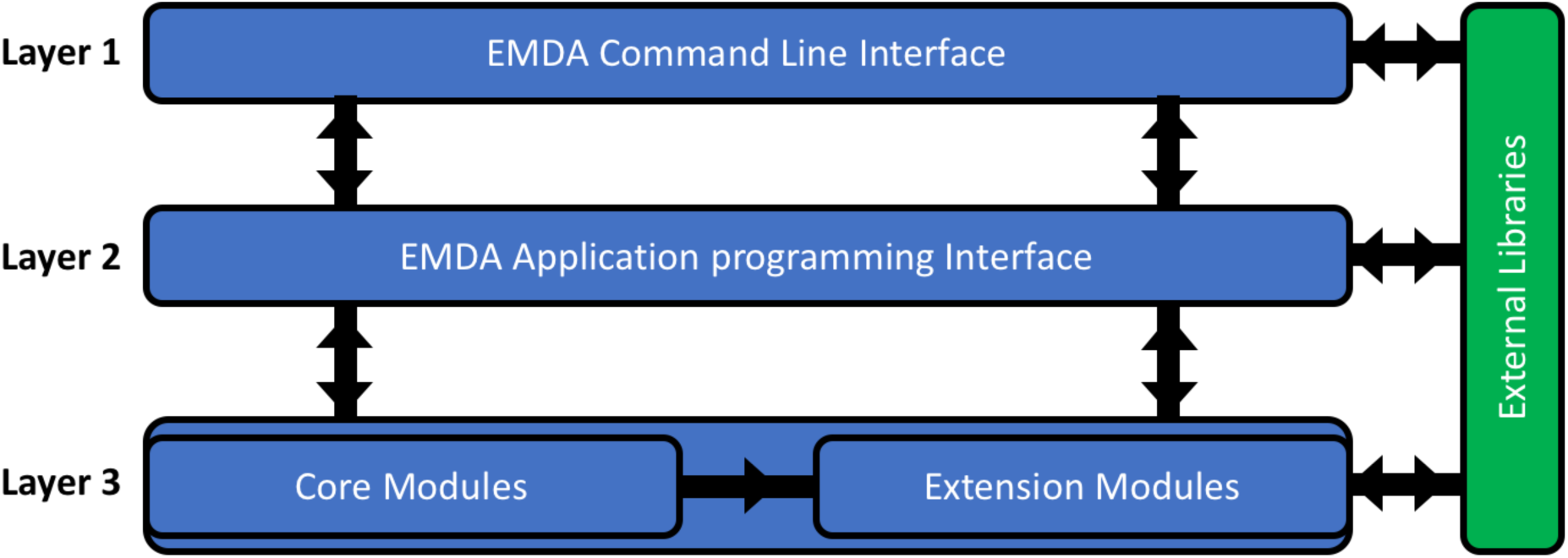
Architecture of EMDA library. The three Python code layers are shown in blue and the layer of external libraries is shown in green. The black arrows show the data flow and the functional dependencies.

EMDA uses open-source, standalone Python library *mrcfile* (Burnley et al., 2017) for the reading, writing and validating EM files in the standard MRC2014 format (Cheng et al., 2015). Also, EMDA uses *gemmi* (https://gemmi.readthedocs.io) for reading and writing atomic coordinate files, and ProSHADE (Nicholls et al., 2018; Tykac, 2018) for symmetry detection in EM maps.

## 3. Methods

In this section we outline the mathematical framework for local correlation and probability-based methods in EMDA. The notations we use throughout this text and in the appendices are summarised in Table 1.

**Table 1:** Table of notation

| Notation | Description |
| --- | --- |
| fullmap | Map obtained by averaging half data reconstructed maps |
| $cov(X, Y)$ | Covariance between random variables $X$ and $Y$ |
| $var(X)$ | Variance of the random variable $X$ |
| $x$ and $s$ | 3D column vectors in real and Fourier space |
| $m(x)$ | Convolution kernel |
| $\psi_i(x)$ | Cryo-EM map number $i$ |
| $var_{i,m}(\psi_i(x))$ | Local variance of $\psi_i(x)$ calculated with the kernel $m(x)$ |
| $cov_{12,m}(\psi_1(x)\psi_2(x))$ | Local covariance between $\psi_1(x)$ and $\psi_2(x)$ calculated with the kernel $m(x)$ |
| $CC_{12,m}(x) = \frac{cov_{12,m}(\psi_1(x)\psi_2(x))}{\sqrt{var_{1,m}(\psi_1(x))var_{2,m}(\psi_2(x))}}$ | Local correlation coefficient calculated between $\psi_1(x)$ and $\psi_2(x)$ with the kernel $m(x)$ |
| $CC_{half,m}(x)$ | Local correlation calculated between halfmaps |
| $CC_{full,m}(x) = \frac{2CC_{half,m}(x)}{1 + CC_{half,m}(x)}$ | Local correlation in the fullmap (Rosenthal and Henderson, 2003) |
| $CC_{map,model,m}(x)$ | Local correlation calculated between the fullmap and the atomic model-based map |
| $\mathcal{N}_2\left(F_{o,j}; k_j F_{t,j}, \frac{1}{2}\sigma_{n,j}^2\right) = \frac{1}{\pi\sigma_{n,j}^2} e^{-\frac{\vert F_{o,j} - k_j F_{t,j}\vert ^2}{\sigma_{n,j}^2}}$ | Two-dimensional Gaussian distribution with mean $k_j F_{t,j}$ and variance $\frac{1}{2}\sigma_{n,j}^2$ of $j^{\text{th}}$ map. Since there is no correlation between real and imaginary parts, a single variance is used to describe this distribution. |
| $\mathcal{N}_{2N}\left(\mathbf{F}_o; \mathbf{kF}_t, \frac{1}{2}\mathbf{\Sigma}\right) = \frac{1}{\pi^N \det(\mathbf{\Sigma})} e^{-(\mathbf{F}_o - \mathbf{kF}_t)^T \mathbf{\Sigma}^{-1} (\mathbf{F}_o - \mathbf{kF}_t)^*}$ | $2N$ dimensional Gaussian distribution with mean $\mathbf{kF}_t$ and covariance $\frac{1}{2}\mathbf{\Sigma}$. Since there is no correlation between real and imaginary parts $N \times N$ covariance matrix is used to describe the $2N$ dimensional random variable*. |
| $\mathbf{F}_o(s) = (F_{o,1}(s), F_{o,2}(s), \dots, F_{o,N}(s))$ | A column vector formed by complex Fourier coefficients of observed maps |
| $\mathbf{F}_t(s) = (F_{t,1}(s), F_{t,2}(s), \dots, F_{t,N}(s))$ | A column vector formed by complex Fourier coefficients of unknown “true” maps |
| $\mathbf{F}_c(s) = (F_{c,1}(s), F_{c,2}(s), \dots, F_{c,N}(s))$ | A column vector formed by complex Fourier coefficients calculated from models |
| $\mathbf{E}_o(s) = (E_{o,1}(s), E_{o,2}(s), \dots, E_{o,N}(s))$ | A column vector formed by normalized complex Fourier coefficients of observed maps |
| $E_{o,j}(s) = \frac{F_{o,j}(s)}{\sqrt{\Sigma_{o,s,jj}(s) + \sigma_j^2(s)}}$ | Normalized complex Fourier coefficients of $j^{\text{th}}$ map. |
| $R_j$ and $t_j$ | Transformation matrix and translational vector in 3D to be applied for the map number $j$ |
| $\mathbf{F}(\mathbf{R}s)e^{2\pi i \mathbf{s}^T \mathbf{t}} = (F_1(R_1 s)e^{2\pi i s^T t_1}, \dots, F_N(R_N s)e^{2\pi i s^T t_N})$ | $N$ dimensional column vector of complex Fourier coefficients after application of transformation. Usually, but not necessarily, $(R_1 = I, t_1 = 0)$ is the identity transformation. |
| $\mathbf{D} = \text{diag}(D_1, \dots, D_N)$ | Diagonal matrix formed by scale factors between the true and calculated Fourier coefficients |
| $\mathbf{k} = \text{diag}(k_1, \dots, k_N)$ | Diagonal matrix formed by blurring parameters |
| $\Sigma$ | $N \times N$ covariance matrix of “true” maps without blurring |
| $\Sigma_{o,s} = \mathbf{k}^T \Sigma \mathbf{k} = \mathbf{k} \Sigma \mathbf{k}$ | $N \times N$ covariance matrix of signals calculated using observed maps |
| $\sigma^2 = \text{diag}(\sigma_{o,1}^2, \sigma_{o,2}^2, \dots, \sigma_{o,N}^2)$ | Diagonal matrix formed by variances of observational noise |
| $\langle \mathbf{F}_t \rangle = (\langle F_{t,1} \rangle, \langle F_{t,2} \rangle, \dots, \langle F_{t,N} \rangle)$ | A column vector formed by the expectation values of the true maps |
| $fsc = 2fsc_{half}/(1 + fsc_{half})$ | FSC in the fullmap converted from halfmap FSC (Rosenthal and Henderson, 2003) |
| $\sqrt{fsc} = \text{diag}(\sqrt{fsc_1}, \sqrt{fsc_2}, \dots, \sqrt{fsc_N})$ | Diagonal matrix formed by square root of fullmap FSC values. It is also an estimate for FSC between fullmap and “true” signal (Rosenthal and Henderson, 2003) |
| $\rho_s$ | $N \times N$ correlation matrix between true maps |
| $\rho_{s,ij} = \frac{\Sigma_{ij}}{\sqrt{\Sigma_{ii}\Sigma_{jj}}}$ | Correlation coefficient between true maps $i$ and $j$. An element in $\rho_s$. |
| $\rho_o$ | $N \times N$ correlation matrix between observed maps |
| $\rho_{o,ij} = \frac{\Sigma_{o,s,ij}}{\sqrt{(\Sigma_{o,s,ii} + \sigma_i^2)(\Sigma_{o,s,jj} + \sigma_j^2)}}$ | Correlation coefficient between observed maps $i$ and $j$. An element in $\rho_o$. |

\* It should be noted that the covariance matrix is $\Sigma \otimes \mathbf{I}_2$ , i.e. Kronecker product of $N \times N$ covariance and 2-dimensional identity matrices. The reason of such covariance matrix structure is that there are not correlations between real and imaginary parts of Fourier coefficients and the variance of real and imaginary parts of each Fourier coefficient are equal to each other.

### 3.1. Local correlation in real space

Pearson’s product-moment sample correlation coefficient (CC) has been extensively used for various purposes in X-ray crystallography (Karplus and Diederichs, 2012; Tickle, 2012) and in cryo-EM (Van Heel, 1987). The CC depends on the signal and noise levels. If we assume that the noise variance is constant within the masked map then for a given data the CC will be an indicator of the signal in the data. Care should be exercised in its interpretation as any systematic behaviour will be considered as signal. Since the CC is calculated using the data, its variance depends on the volume of the data being used.

Local CC in real space can be calculated using the formula for the weighted Pearson’s product moment sample correlation coefficient with weights equal to the kernel. The local CC for two maps *Ψ*_1_*(x)* and *Ψ*_2_*(x)* is:

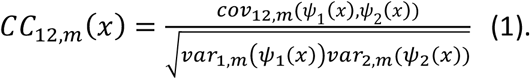

*Cov*_12,*m*_ (*Ψ*_1_(*x*),*Ψ*_2_(*x*)) and *var*_*i,m*_(*Ψ*_*i*_(*x*)) are the local covariance and variances for *Ψ*_1_*(x)* and *Ψ*_2_*(x)* calculated with a kernel *m*(*x*). The kernel is normalized such that 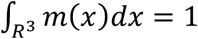.

The expression for local covariance is,

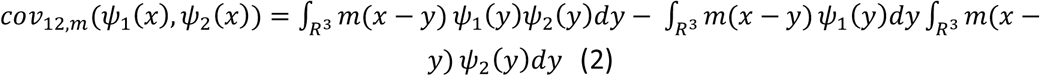

Similar expressions can be written for local variances *var*_*i,m*_(*Ψ*_*i*_(*x*)). Note that the eq. 2 can be readily evaluated using the convolution theorem (see Appendix A).

Such correlations could be calculated for any pairs of maps. When it is calculated using half maps (*CC*_*half,m*_(*x*)) reconstructed from randomly chosen half of the particles, then it indicates the local signal to noise ratio, whereas the local correlation between observed and calculated maps (*CC*_*map,model,m*_(*x*)) indicates local agreement between atomic model and observed map. Similarly, the local correlation between two different observed maps indicates local common signals between them.

The correlation calculated using half maps is converted to that of fullmap using the following formula (Rosenthal and Henderson, 2003)

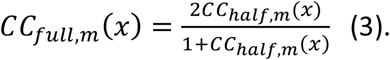

In the local correlation calculation, EMDA uses a spherically symmetric kernel defined as:

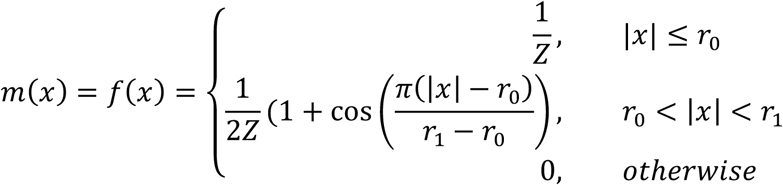

Where *r*_*0*_ and *r*_1_ are the radii of inner and outer concentric spheres. *Z* is the coefficient that makes sure that the total integral of *m(x)* is equal to 1.

The size of the kernel (i.e. *r*_1_) should be chosen such that the number of data points included is sufficient to calculate a reliable statistic. Both too small or too large kernels lead to inaccurate correlations due to insufficient data points or loss of locality, respectively. In the current implementation of EMDA, *r*_1_ should be chosen by trial-and-error. Both *r*_*0*_ and *r*_1_ are in pixel unit and by default 2 pixels are used to soften the edge of the mask, in other words *r*_1_ *= r*_*0*_ + 2.

The *CC*_*full,m*_*(x)* (hereafter *CC*_*full*_) depends on the local signal strength. It has two implications: 1) it depends on the local variation of the signal, and hence different parts of the map with different mobilities will have differing correlations; 2) it will depend on global sharpening/blurring parameters, i.e. maps sharpened with different B values will have different local *CC*_*full*_.

It should also be noted that the *CC*_*map,model,m*_*(x)* (hereafter *CC*_*map,model*_) calculated between an atomic model and a given map (fullmap) depends on atomic coordinates, occupancies and B values. Therefore, to obtain the best possible *CC*_*map,model*_ it should be calculated using a refined model with optimized atomic B values.

In order to compare *CC*_*map,model*_ with *CC*_*full*_ all maps should be weighted appropriately. To achieve this, Fourier coefficients of all maps are normalized and weighted by FSC in resolution bins. All correlation examples discussed in this paper used such normalized and weighted maps. The details are in Appendix D.

Let us assume that errors in the observations are additive and they follow a Gaussian distribution with zero mean. Also, assume that there is no correlation between the noise and the calculated map from the atomic model. This is true only when there’s no overfitting. Under these assumptions, a relationship between *CC*_*map,model*_ and *CC*_*full*_ useful for validation is given by eq. 4. The full derivation of eq. 4 is given in Appendix A as well as in (Nicholls et al., 2018).

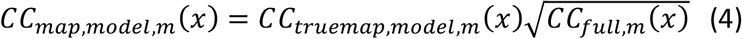

According to eq. 4, the *CC*_*map,model*_ *(x)* is equal to 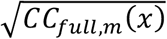 only when *CC*_*truemap,model,m*_ *(x) =* 1, i.e. perfect model. Since this situation is almost never realised, the *CC*_*map,model,m*_*(x)* should always be less than 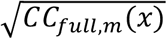. If *CC*_*map,model,m*_*(x)* is greater than 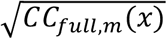, that could be an indication of overfitting.

### 3.2. Parameter estimation and map calculation: likelihood and posterior distribution

As in any application of Bayesian computations to the data analysis we need two probability distributions: 1) probability distribution of observations given parameters to be estimated – likelihood function and 2) probability distribution of unknown signal given observations and current model parameters – posterior probability distribution. The details are given in Appendix C.

#### Likelihood function

The negative log likelihood function in the absence of atomic models and in the presence of multiple related maps is (see Appendix C for details):

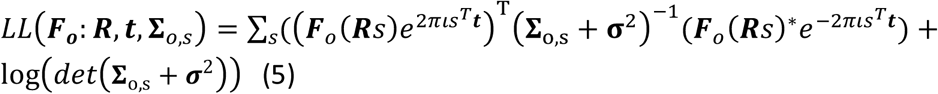

Where ***F***_***o***_(*s*) is a vector of Fourier coefficients of the observed maps, ***R*** and ***t*** are the vectors formed by rotation and translation parameters for each map, respectively, **Σ**_o,s_ is the covariance matrix between “true” maps calculated using observed maps and ***σ***^2^ is a diagonal matrix of noise variances. In EMDA, the above likelihood function is implemented to estimate parameters between a pair of maps where one map is static the other moves onto it. In the case of estimation of transformation parameters, the only terms that depend on adjustable parameters are the cross-terms in the eq. 5:

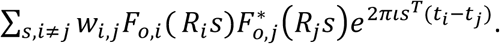

*F*_*o,i*_ and *F*_*o,j*_ are the Fourier transforms of *i*^th^ and *j*^th^ maps, *R*_*j*_ and *t*_*j*_ are the rotation and translation parameters. *W*_*i,j*_ is related to the corresponding term in the inverse of the covariance matrix and is related to FSC between maps. For parameter estimation **Σ**_o,s_ and ***σ***^2^ do not need to be estimated separately, their sum is used in eq. 5.

Note that if we relax the conditions that ***R*** is a rotation matrix, the same formula also allows refining the magnification parameters. In cryo-EM, we assume that the magnification is a scalar parameter and ***R*** becomes a diagonal matrix with the same magnification parameter in magnification-only-refinements. The covariance matrices and transformation parameters are estimated iteratively.

The algorithm in EMDA for transformation estimation includes following steps. 1) starting with initial rotation and translation parameters the covariances are calculated and converted them into weights to calculate the functional value. 2) the derivatives of translation and rotation are calculated and the shifts are estimated. 3) the current translation and rotation are updated and applied on maps. The covariances are recalculated and the new functional value is evaluated. 5) the new functional value is compared with that of the previous iteration, and if the convergence criterion is met the final maps are output and the transformation is retained. Otherwise, the process continues at step 2 with the next cycle of iteration.

#### Posterior distribution

For map calculations we need the probability distribution of the unobserved “true” maps given the current state of atomic models as well as observations. In the absence of atomic models this distribution is a multivariate Gaussian with mean (see Appendix C for details):

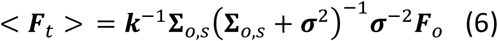

where ***k*** is the diagonal matrix formed by the blurring parameters, ***σ***^2^ is the diagonal matrix formed with the variance of the noise in the observations, **Σ**_*o,s*_ is the covariance matrix between “true” maps calculated using observed maps, ***F***_*o*_ is a vector of Fourier coefficients of observed maps, < ***F***_*t*_ > is a vector of expectation values of the “true” map Fourier coefficients.

Since ***k*** is unknown, we replace it with the standard deviation of the signal as explained in (Yamashita et al., 2021). After some algebraic manipulation we get (see Appendix C):

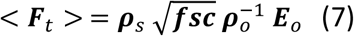

where ***ρ***_*s*_ is the correlation matrix between Fourier coefficients of “true” maps, ***ρ***_*o*_ is the correlation matrix between Fourier coefficients of observed maps, ***E***_*o*_ is a vector of normalised Fourier coefficients of observed maps and < ***F***_*t*_ > is a vector of the expectation values of the normalised Fourier coefficients of the “true” maps.

Note that if we know the blurring factor, it might be better to use them in the map calculation. However, observations alone do not allow us to calculate these quantities, and they need to be estimated using different methods.

## 4. Results and Discussion

In this section, we demonstrate the use of local correlation, map overlay and magnification refinement implemented in EMDA package through examples.

### 4.1. Examples of use of local correlation

#### 4.1.1. Model-map differences by local correlation

To demonstrate the use of local correlation to detect model-map differences, we used archaeal 20S proteasome (EMD-5623) map with overall resolution 3.3 Å and the corresponding atomic model 3j9i (Li et al., 2013). The atomic model was refined against the fullmap to a resolution of 3.3 Å using REFMAC5 (Nicholls et al., 2018). Using the refined model, an EM map to 3.3 A was computed in EMDA using *gemmi* (https://gemmi.readthedocs.io). Local correlations were calculated within a kernel of radius *r*_*1*_ =4 pixels (pixel size = 1.22 Å). The *CC*_*full*_ was calculated using the normalized and weighted halfmaps (see Appendix A). Similarly, the *CC*_*map,model*,_ was calculated between the normalized and weighted fullmap and the normalized and weighted calculated map. Fig. 2a shows the primary map density near residues Lys52-Val54 of chain U of the 3j9i model coloured by the 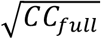 and the superimposed model. One can appreciate a moderate signal, but the model is outside the density. The same density coloured by *CC*_*map,model*_ (Fig. 2b) shows a low correlation resulting from the misplaced model. Fig. 2c shows the corresponding part of the refined atomic model coloured by *CC*_*map,model*_ and it also highlights those residues with low correlation. Thus, *CC*_*map,model*_ can highlight areas with map-model discrepancies, while 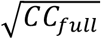 can be used to validate the existence of a signal. A comparison of *CC*_*map,model*_ versus 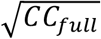 is useful not only to pinpoint map-model differences, but also to identify viable ways to minimise them. Moreover, colouring the atomic coordinates by *CC*_*map,model*_ is an effective way to identify misplaced regions in the model.

**Fig. 2.**
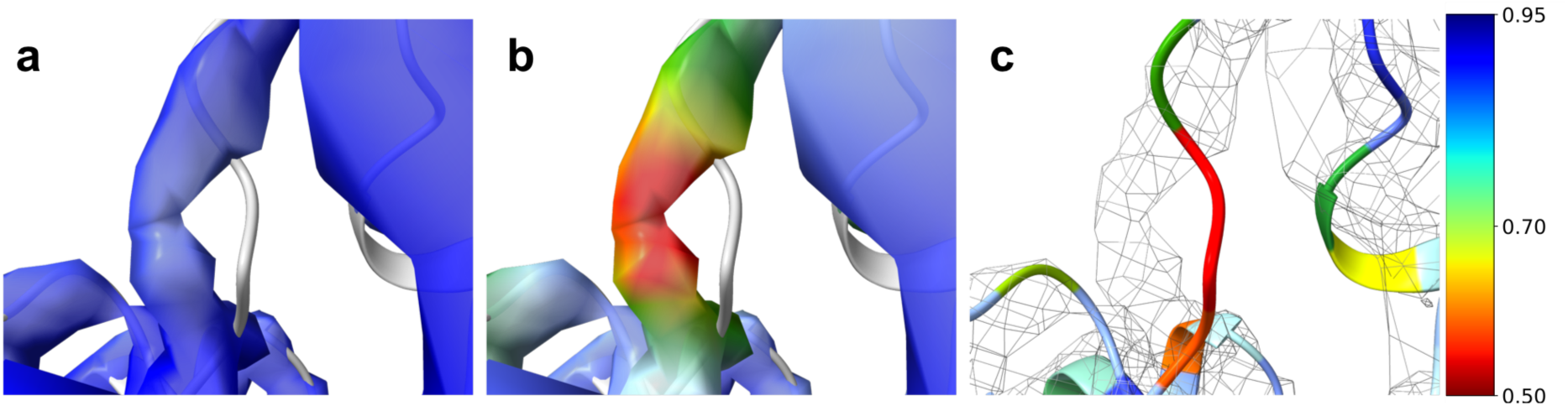
Identifying model-map discrepancies by local correlation. a) EMD-5623 primary map density near residues Lys52-Val54 of chain U of the 3j9i model coloured by 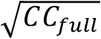. b) the same density coloured by *CC*_*map,model*_. c) corresponding atomic model coloured by *CC*_*map,model*_ showing low correlation residues Lys52-Val54 of chain U. The figure was made with ChimeraX (Pettersen et al., 2021).

#### 4.1.2. Unmodeled regions by local correlation

Next, we used SARS-CoV-2 spike protein structure EMD-11203 and the corresponding model 6zge (Wrobel et al., 2020) to demonstrate the use of local correlation to highlight an unmodeled density. This density has been modelled as linoleic acid (LA) in the homology model 6z5d (Toelzer et al., 2020).

First, we present the use of 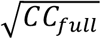 and *CC*_*map,model*_ to identify the unmodeled density in the map. Then, we compare those local correlations against the local correlation calculated from the model with the ligand.

Using the normalized and weighted EMD-11203 halfmaps the *CC*_*full*_ was calculated within a kernel of radius *r*_*1*_ =3 pixels (pixel size = 1.087 Å). The size of the kernel was chosen to maximize the variation of local correlation and minimize the leakage of correlation from the surrounding (a comparison of correlations calculated using different kernel sizes is given in Supplementary materials). The model 6zge was refined against the EMD-11203 fullmap using REFMAC5 (Nicholls et al., 2018) to optimise atomic coordinates and the B values. Using the refined model an EM map was computed to 2.6 Å in EMDA using *gemmi* (https://gemmi.readthedocs.io). The *CC*_*map,model*_ was calculated using the normalized and weighted EMD-11203 fullmap and the normalized and weighted model-based map. EMD-11203 primary map was coloured by 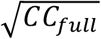 and *CC*_*map,model*_, and their comparison highlighted an unmodeled structured densities located near all receptor binding domains. One such density coloured by 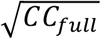 and *CC*_*map,model*_ are shown in Fig. 3a and 3b, respectively. The high 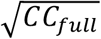 implies the density is real, but low *CC*_*map,model*_ implies there is no corresponding model. Next, the homology model 6zb5 with LA was fitted on the EMD-11203 fullmap and refined using REFMAC5 (Nicholls et al., 2018), and the *CC*_*map,model*_ was recalculated. The improved *CC*_*map,model*_ for the ligand region is shown in Fig. 3c and this improvement is due to the presence of LA in the model.

**Fig. 3.**
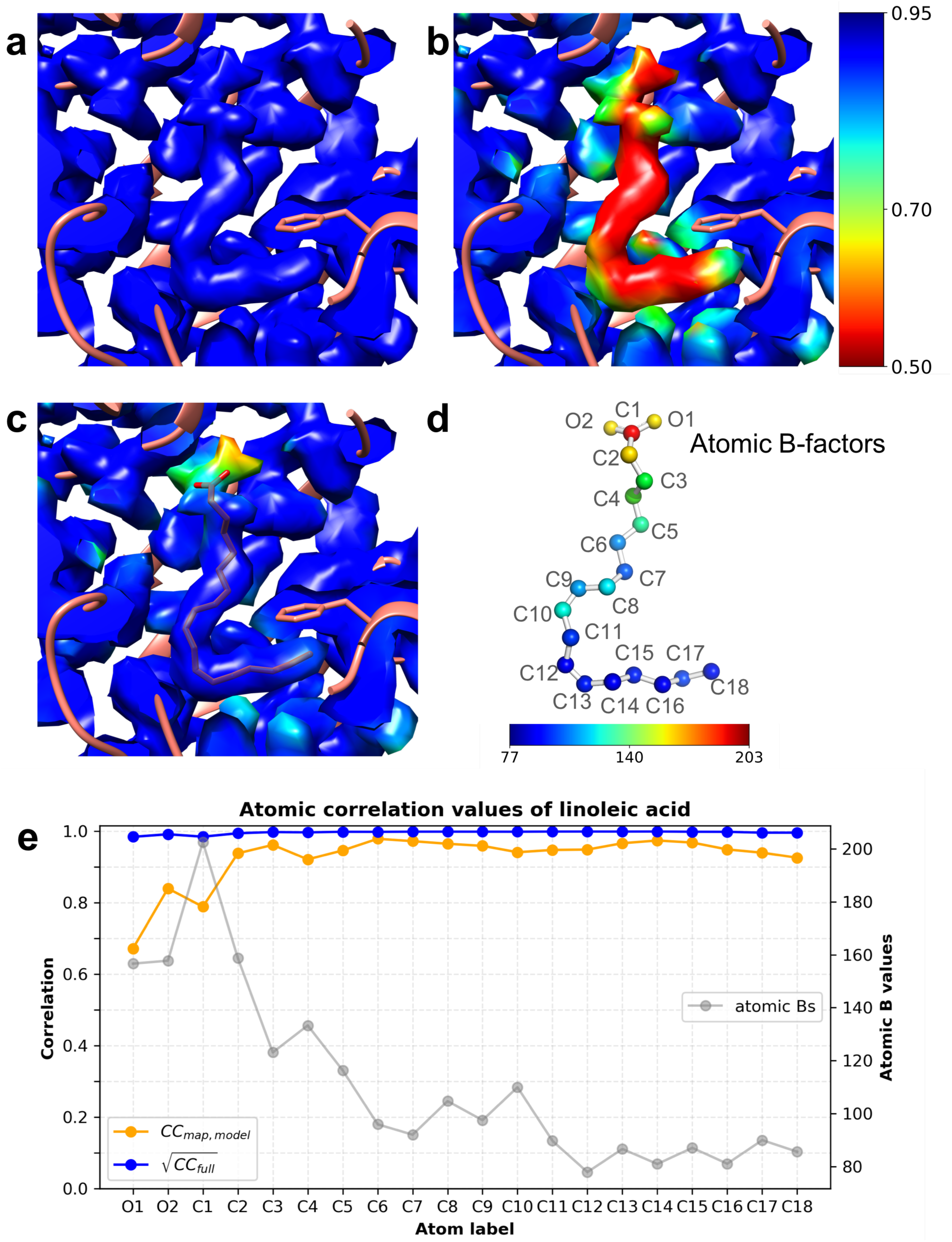
Use of local correlation to identify unmodeled linoleic acid (LA) in EMD-11203 map. a) unmodeled ligand density in the primary map coloured by the 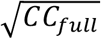. High correlation indicates the presence of a strong signal. b) the same density coloured by *CC*_*map,model*_ calculated between the fullmap and the refined 6zge model using normalized and weighted densities. The correlation in this region is low compared to its surrounding. c) ligand density coloured by *CC*_*map,model*_ calculated between the fullmap and the refined model with LA using normalized and weighted densities. Densities in a, b and c panels were contoured at the same level. Those figures were made with Chimera (Pettersen et al., 2004). d) distribution of atomic B values of refined LA where the atoms are coloured by the B values. This figure was made with PyMOL (Schrödinger, 2020). e) distribution of atomic correlation values at refined LA coordinates. *CC*_*map,model*_ and 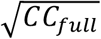 are shown with orange and blue, respectively. Also, the atomic B values are shown in grey.

Fig. 3d shows the refined LA molecule whose atoms are coloured according to the atomic B values. The overall trend shows that B values in the hydrophobic tail are relatively small, but they gradually increase towards the hydrophilic carboxyl group. In Fig. 3e, atomic 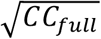 and *CC*_*map,model*_ are plotted in blue and orange, respectively, along with the atomic B values in grey. The atomic correlation values were obtained from correlation maps by interpolating at atomic positions.

The 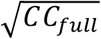 is close to 1 throughout the molecule, but largest variation is seen in the carboxyl group. The *CC*_*map,model*_ is lower than the 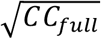 in all atoms, and its variation is larger than that of 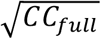. The difference in atomic *CC*_*map,model*_ for carbonyl O1 and O2 is significant despite their similar B values. The carbonyl group is anchored by the neighbouring Arg408 and Gln409 residues through H-bonding with O2 atom (Toelzer et al., 2020, Fig. 3c), but O1 atom does not seem to have close neighbours thus its atomic correlation may be compromised by the surrounding noise. It should also be noted that at some atoms there is a leakage of correlation. This effect is pronounced at atoms O2, C10, C14 and C18 in Fig. 3b.

A comparison of 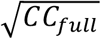 versus *CC*_*map,model*_ can highlight map-model differences as shown by the previous examples. In model building and refinement, we aim at explaining the full signal in the map by the model, and hence the chance of building model into the noise is unavoidable. In such situations, the local correlation can be a helpful tool to monitor the overfitting. According to eq. 4 the inequality 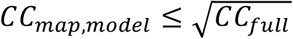 should hold when there is no overfitting, and Fig. 3e shows such a situation.

### 4.2. Examples of use of map overlay

#### 4.2.1. EMDA map overlay

To demonstrate the overlay method, we have used cryo-EM maps EMD-21997 and EMD-21999 (Henderson et al., 2020) of SARS-CoV-2 spike protein, whose resolutions are 2.7 Å and 3.3 Å, respectively. The former map is in rS2d locked state in which all three receptor binding domains (RBDs) are down and locked, thus maintaining C3 symmetry. Whereas the latter map is in u1S2q state in which one of the RBDs is open causing the whole structure to be in C1. In this example, we estimate the movement of one of the down RBDs in EMD-21999 map relative to one of the down RBDs in EMD-21997 map using EMDA overlay operation. We kept the primary EMD-21997 map and the corresponding 6×29 atomic model static, while the primary EMD-21999 map and the corresponding 6×2a atomic model moving. First, we overlaid EMD-21999 map (Fig. 4a(ii)) on EMD-21997 map (Fig. 4a(i)) and the resulting transformation (relative rotation = 8.35°, translation = 4.14 Å) was applied on the 6×2a atomic model. The overlaid maps are shown in Fig. 4a(iii), and they are the starting maps for the subsequent domain overlay (shown by 4a(iv) and (v) for static and moving maps, respectively). Next, a pair of RBDs located proximity to each other on 4a(iii) were extracted within model generated masks. The extracted RBDs are shown in 4a(vi) and 4a(vii) and their superposition before fit optimisation is shown in 4a(viii). Next, their relative fit was optimized and at the convergence the relative rotation and translation values were 3.38° and 1.76 Å, respectively. The superposed domains after the fit optimisation is shown in 4a(ix). These rotation and translation values indicate the movement of the selected RBD of EMD-21999 map relative to the corresponding RBD of EMD-21997 map in the same coordinate frame. Finally, the estimated transformation between domains was applied on the 6×2a RBD coordinates to bring it on the static model (6×29). Fig. 4a(x) and (xi) present the superposition of models before and after the transformation has been applied, respectively. Fig. 4b shows the FSC curves calculated between the two domains before and after the overlay optimisation.

**Fig. 4.**
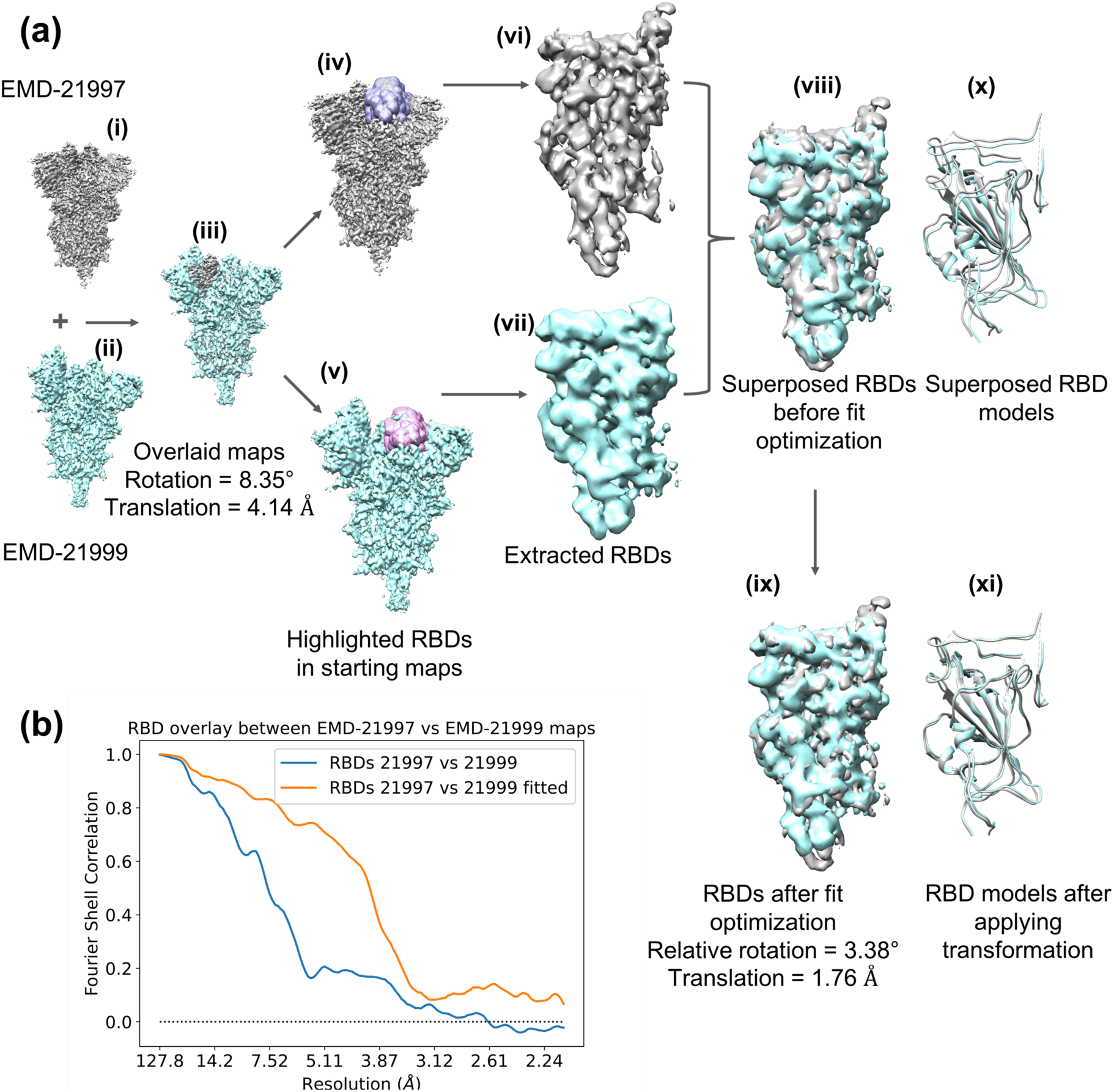
Map superposition in EMDA illustrated using EMD-21997 and EMD-21999 maps. (a) keeping EMD-21997 map (i) static, EMD-21999 map (ii) was moved to obtain the optimal overlay between them (iii). Starting from the overlaid maps (iv) and (v), RBDs were extracted using masks. The extracted RBDs (vi) and (vii) were superposed (viii) and optimized their overlay (ix) in EMDA. The final values of relative rotation and translation are 3.38° and 1.76 Å, respectively. The same transformation was applied on the model 6×2a of the moving map. The superposition of 6×29 (static, grey) and 6×2a (moving, cyan) RBD models before (x) and after (xi) the domain transformation being applied. This figure was made with Chimera (Pettersen et al., 2004). (b) FSC between static and moving RBDs before (blue) and after (orange) the overlay optimization.

#### 4.2.2. EMDA magnification refinement

Magnification refinement in EMDA involves 1) resampling the target map on the reference grid to make sure both maps refer to the same coordinate system, 2) superposition of the target map on the reference and refining the magnification of the target map, iteratively. To demonstrate the magnification refinement in EMDA, we intentionally introduced a −5% magnification error in one of the half maps of Haemoglobin (half1 of EMD-3651; (Khoshouei et al., 2017)) to yield the initial map, and let EMDA to refine its magnification against the half1 map (original map). The pixel sizes of the original and the magnification modified maps (initial map, Fig. 5a) are 1.05 and 0.998 Å, respectively. EMDA optimized the magnification of the initial map relative to the original map to yield the magnification adjusted map (Fig. 5a) with the pixel size 1.05 Å. Fig. 5b shows the FSC curves for the initial and the adjusted maps calculated against the original map. The increase from initial map to adjusted map is due to the correction in the magnification. To validate the accuracy of refinement, the FSC for adjusted map is compared with the half data FSC (Fig. 5c) and they are in very good agreement.

**Fig. 5.**
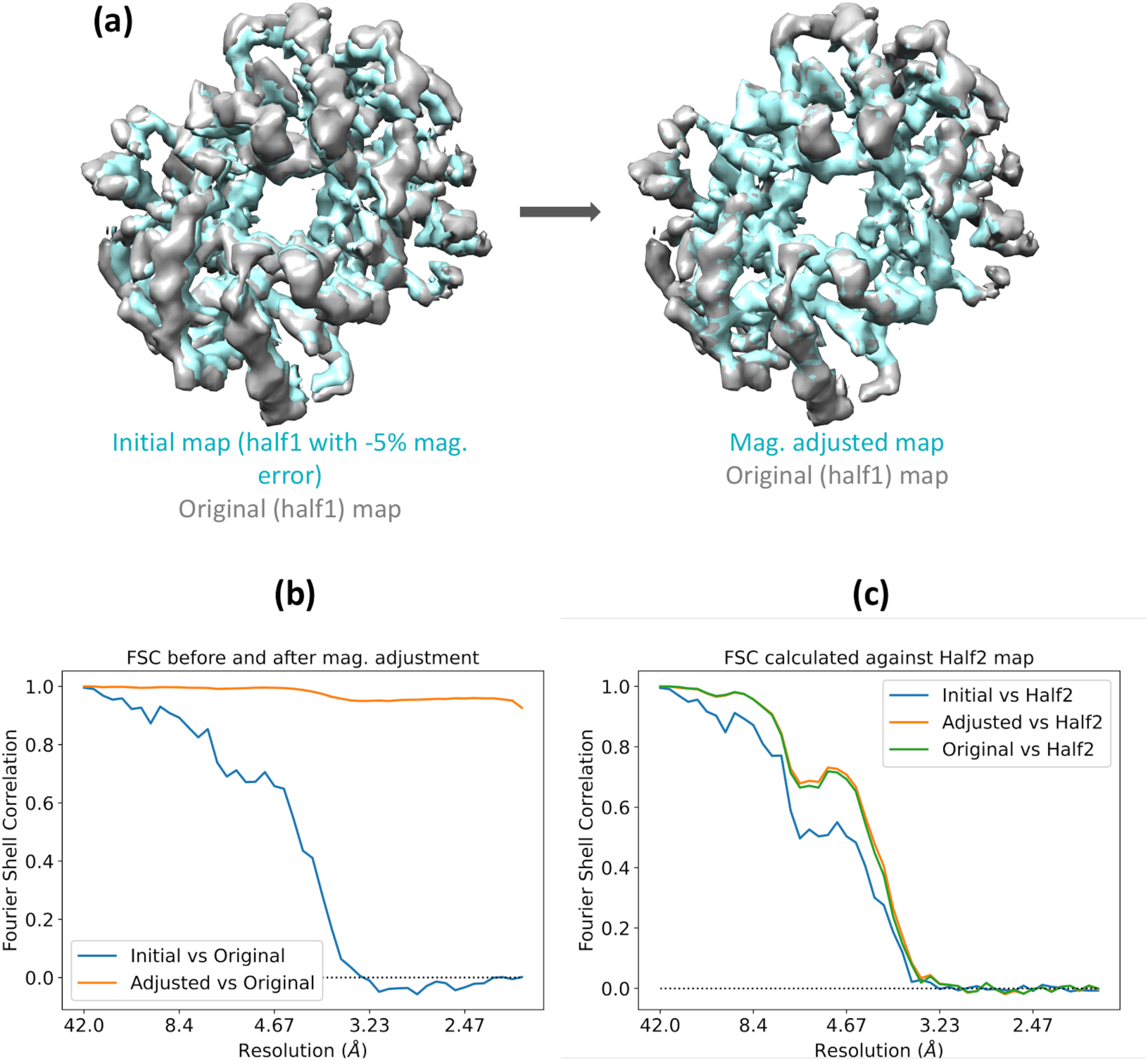
The magnification refinement in EMDA using Haemoglobin data (EMD-3651). (a) the superposition of the original (half1) map (in grey) on the initial map (in cyan) obtained by introducing a −5% magnification error on the original map is improved after magnification correction (adjusted map shown in cyan). This figure was made with Chimera (Pettersen et al., 2004). (b) FSC between initial and adjusted maps against the original map indicating improvement in the superposition due to correction in magnification. (c) FSC curves for initial and adjusted maps calculated against the half2 are shown in blue and orange, respectively. The increase of FSC from blue to orange is due to the improved magnification. The green curve is the FSC between the half maps and it serves as the ground truth.

In the next example, we illustrate the estimation of the relative magnification differences of two cryo-EM maps of beta-galactosidase [EMD-7770 (Bartesaghi et al., 2018) and EMD-10574 (Saur et al., 2020)] relative to an X-ray crystallography model 3dyp (Juers et al., 2009). The resolution of EMD-7770 and EMD-10574 are 1.9 and 2.2 Å, respectively. The model 3dyp has been derived from X-ray data with resolution 1.75 Å. Both cryo-EM maps and the crystallographic model possess D2 point-group symmetry. Since one of the cryo-EM primary maps (i.e. EMD-10574) has been lowpass filtered, we used fullmaps generated from half maps for both cryo-EM entries in this analysis. First, all non-polymer atoms of 3dyp model were removed and just the polymers were fitted onto EMD-7770 map in Chimera (Pettersen et al., 2004). Then the model-based map was calculated up to 1.9 Å using REFMAC5 (Nicholls et al., 2018) and it was kept as the crystallographic reference for the subsequent magnification analysis. Both the reference map and the EMD-7770 map have the same pixel size 0.637 Å, while EMD-10574 map has 0.68 Å. Thus, the latter map was resampled on the reference to bring all maps on the same coordinate system. Next, a principal component analysis was performed on the variance-covariance matrices of the reference and resampled maps to bring the orientation of the latter approximately matches that of the reference.

Lastly, the fits and the magnifications of EMD-7770 and the resampled EMD-10574 maps were optimized relative to the reference map, iteratively. This resulted in +0.3 % and +1.7 % magnification differences in EMD-7770 and EMD-10574 maps relative to the reference, respectively. Fig. 6a(i) and (ii) show the superpositions of EMD-7770 (yellow) and EMD-10574 (cyan) maps on the reference (grey). Their magnified portions enclosed by red rectangles are shown in Fig. 6b on the left two columns. The yellow density overlaid on the grey density does not show an obvious offset discernible to human eye in both centre or periphery regions. However, the cyan density shows an offset relative to the grey density. Moreover, this offset increases from the centre to periphery; an indication of the magnification problem. Fig. 6a(iii) and (iv) show the magnification corrected EMD-7770 and EMD-10574 maps overlaid on the reference map, respectively. The magnified portions marked by red rectangles are shown in Fig. 6b on the right two columns for centre and periphery regions. Both yellow and cyan densities overlay on grey density, and the offset seen in the cyan density before the correction has now disappeared confirming that EMD-10574 map indeed suffers from magnification problem. Furthermore, Fig. 6a(v) and (vi) present the masked FSC curves for EMD-7770 and EMD-10574, respectively, before (blue) and after (orange) the magnification has been corrected. The increase in FSC, especially in (vi) is attributed to the improved magnification.

**Fig. 6.**
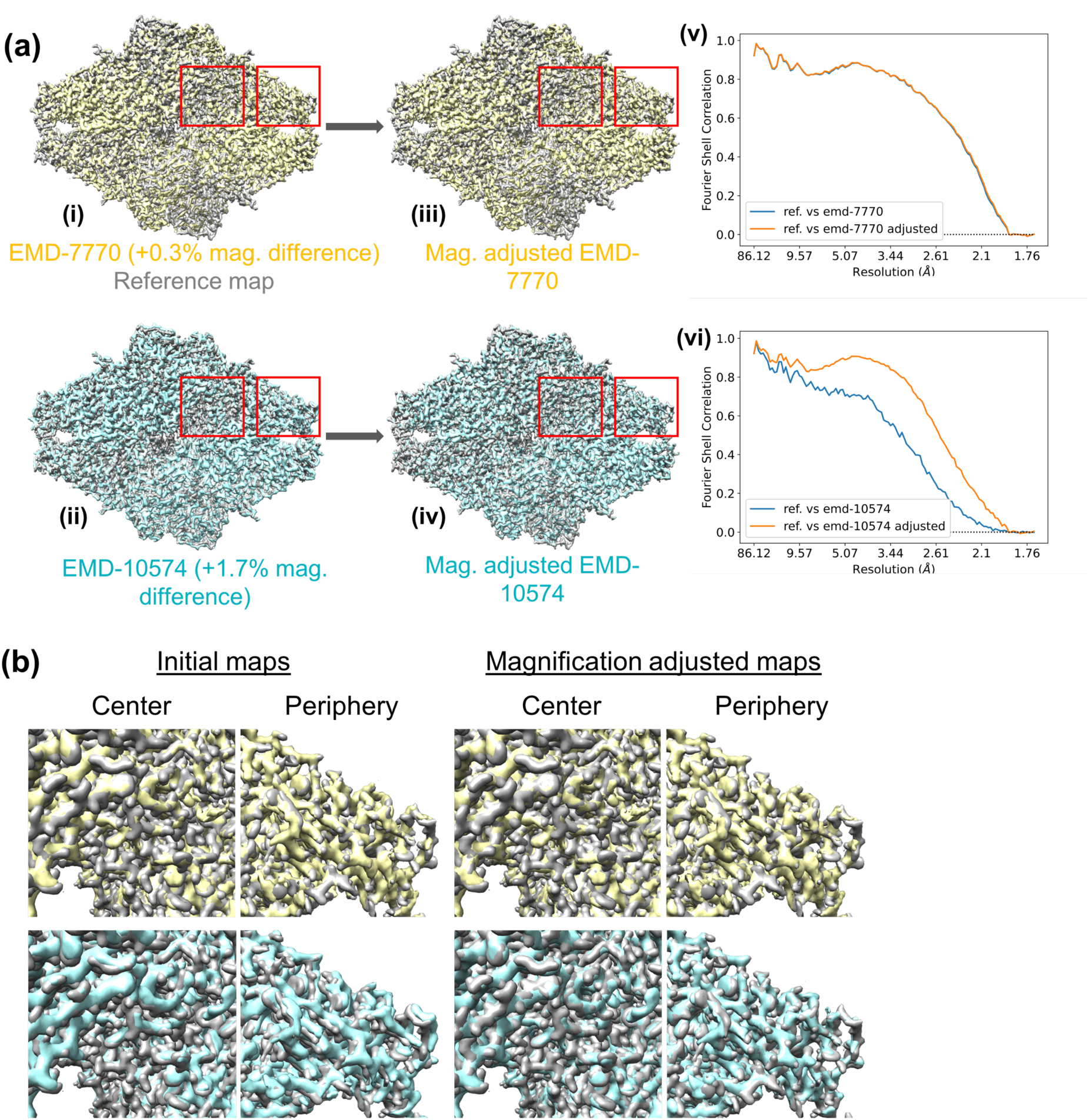
Magnification correction in EMD-7770, EMD-10574 maps relative to the crystallography model 3dyp. (a) the overlaid EMD-7770 (i, yellow) and EMD-10574 (ii, cyan) maps on the reference map (grey) before the magnification optimisation. (iii) and (iv) are the same maps after the optimisation. The magnification differences in EMD-7770 and EMD-10574 relative to the reference are +0.3 % and +1.7 %, respectively. The FSC curves for EMD-7770 and EMD-10574 maps against the reference before and after the magnification adjustment are shown in (v) and (vi), respectively. The blue and orange curves correspond to FSCs before and after the magnification refinement, respectively. The increase in FSC is attributed to the corrected magnification. This figure was made with Chimera (Pettersen et al., 2004). (b) comparison of EMD-7770 map (yellow) and EMD-10574 map (cyan) densities against the reference map (grey) in different regions before and after the magnification correction. See text for details.

Even after the magnification correction, some discrepancies in density overlay were apparent in both EMD-7770 and EMD-10574 maps relative to the reference map. We focused on one monomer unit of EMD-7770 map and extracted it using a model generated mask. The corresponding monomer unit of the reference map was also extracted in similar manner. Fig. 7(i) shows the overlaid EMD-7770 map on the reference after the magnification correction. The monomer units chosen is highlighted within the mask. Extracted monomers are shown in Fig. 7(ii), and one can easily appreciate the rotation of the yellow density relative to the reference grey density due to movements between domains. We estimated the relative transformation between those monomer units and that resulted in 1.02° rotation and 0.12 Å translation (similar analysis was performed using monomers from EMD-10574 map and the reference. That resulted in 0.28° rotation and 0.17 Å translation). Fig. 7(iii) and (iv) show the optimized fit of the monomers and the FSCs between them before (blue) and after (orange) the fit optimisation, respectively. The increase in FSC is attributed to the improved fit.

**Fig. 7.**
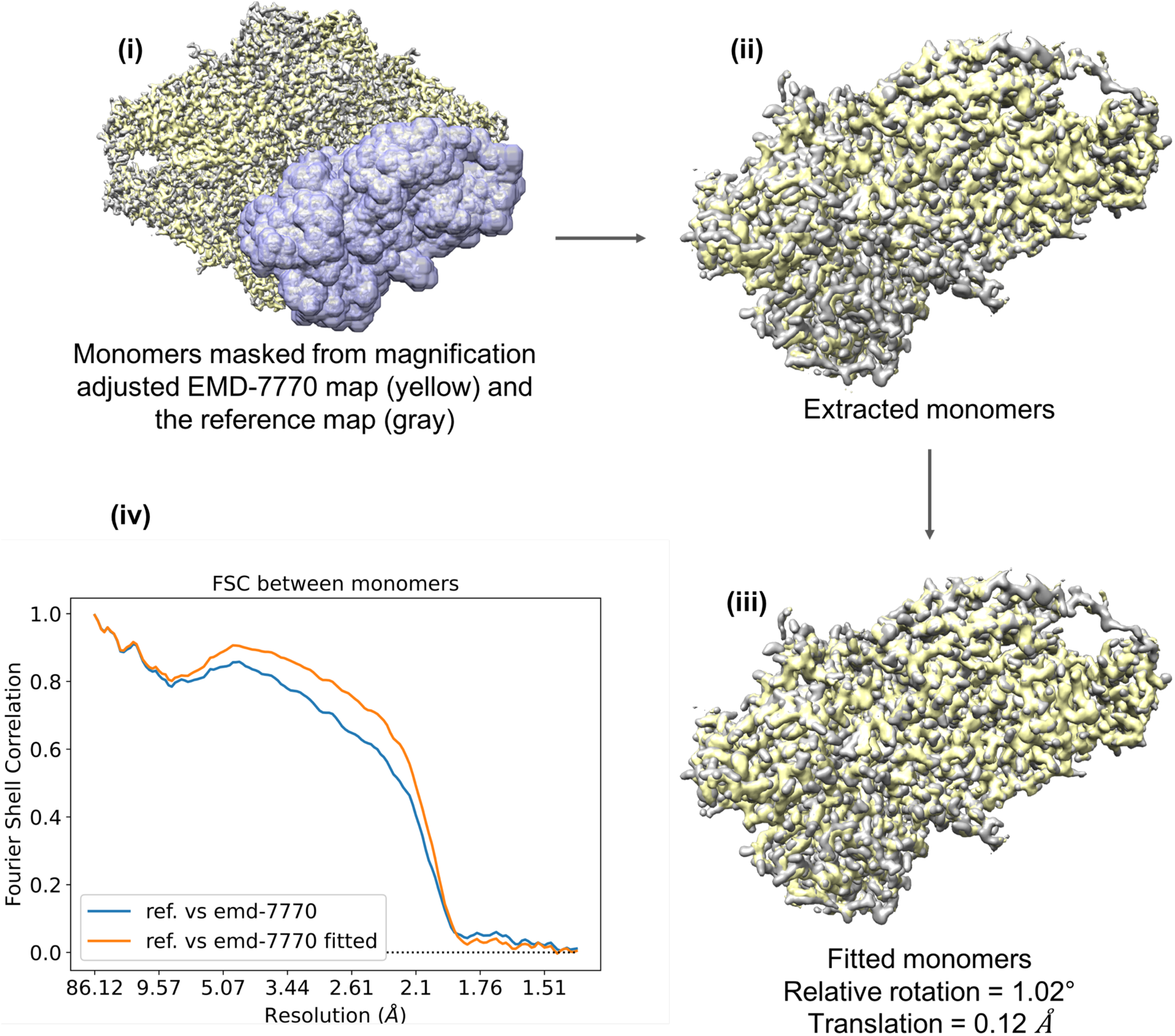
Movement of one monomer unit of EMD-7770 (yellow) relative to the corresponding monomer unit of the reference map (grey). Selected monomers are highlighted in (i) and those extracted are shown in (ii) before the fit optimisation. (iii) the monomer units after the fit optimisation. (iv) FSCs between monomer units before (blue) and after (orange) the fit optimisation. This figure was made with Chimera (Pettersen et al., 2004).

As illustrated in this example, the map magnification is an important factor to consider during structural comparison studies. It should be refined and make sure all structures have the same magnification before comparing for other structural variations. Internal motions such as domain movements should be estimated and compared to other similar structures only if their magnifications are comparable.

## 5. Conclusions

We presented the EMDA Python package to serve the need of map and model validation in cryo-EM. We showed the use of map-model local correlation to identify residues outside the density or those poorly fitted. Since the fullmap local correlation gives an indication of the signal level in the map, it can be used to draw insights about the presence of a signal. Moreover, a comparison of map-model local correlation with fullmap local correlation can be used for validating the model-to-map fit. In one of the examples, we used the local correlation to identify an unmodeled ligand in a map, thereby demonstrating its complementary nature to the difference map. The use of local correlation to identify ligands has the advantage that the correlation naturally offers a way to validate the presence/absence of the density as revealed by the half map local correlation. Also, we showed that correlation values mapped into atoms are useful to study the local signal variations.

Secondly, we presented the likelihood-based map-to-map fitting using an example, where two SARS-CoV-2 structures were first fitted to bring them on the same coordinate frame. Then two receptor binding domains were fitted in the same coordinate frame to estimate their relative movement. The last example illustrated the use of likelihood-based magnification adjustment where the magnifications of two cryo-EM maps relative to an X-ray crystallography derived atomic model have been estimated. The importance of correcting the relative magnification between structures in structure comparison studies have been highlighted.

### Software availability

EMDA is released under the Mozilla Public License Version 2.0 (MPL 2.0) and it is free and open source. The source code is accessible at https://gitlab.com/ccpem/emda. EMDA is distributed as a part of CCP-EM suite and also available via Python Package Installer (`pip`). EMDA’s documentation is available at https://emda.readthedocs.io, and we encourage the reader to look at the documentation for most recent functionalities and up-to-date instructions.

## Acknowledgements

RW was supported by the MRC and the Wellcome Trust (grant numbers: MC_UP_A025_1012, 208398/Z/17/Z), GNM and KY were supported by MC_UP_A025_1012, We thank Takanori Nakane for useful discussions, providing test cases and comments on the initial manuscript, Jake Grimmett and Toby Darling for providing scientific computing resources, UKRI, CCP-EM, UK Cryo-EM validation group, the MRC-LMB for creative environment and the users for their valuable feedback.

## Appendix A

### Local correlation in real space

Let *Ψ(x)* and *m(x)* be two functions where the latter is normalized

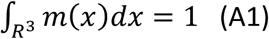

Then the local averages of *Ψ(x)* with the kernel *m(x)* can be written as a convolution operation:

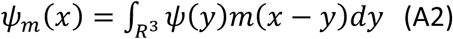

This can be calculated using the convolution theorem:

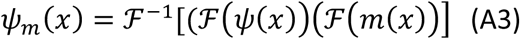

where ℱ is the Fourier transformation operator and ℱ^−1^ is its inverse.

Similarly, the local covariance between *Ψ*_1_*(x)* and *Ψ*_2_*(x)* is given by

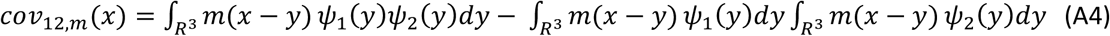

The local correlation between the two functions can be written as

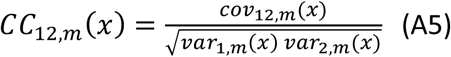

Now, let us assume that there are two noisy maps, each has

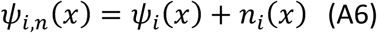

where *Ψ*_*i*_*(x)* and *n*_*i*_*(x)* are the signal and the noise components in *i*^th^ map. If the noise components between the maps, and the noise and signal within as well as between maps are uncorrelated, then the local variance and covariance can be written:

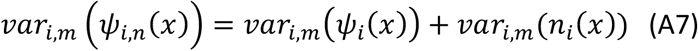

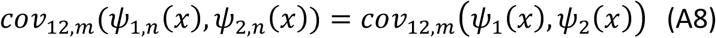

And finally, the local correlation can be written

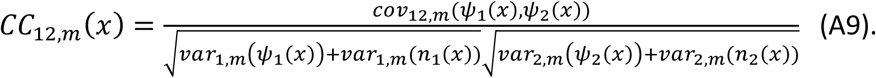

If two maps are cryo-EM half maps, then they share a common signal. In addition, if the noise components have the same variance for both halves then the following relationships hold

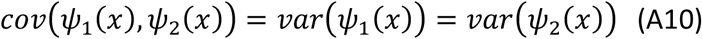

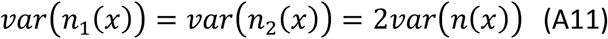

where *var*(*n*(*x*)) is the noise variance in the averaged map.

Thus, the local correlation between half maps is:

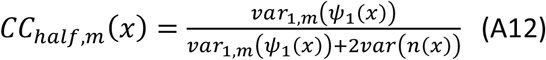

The corresponding local correlation in the full map (Rosenthal and Henderson, 2003) is

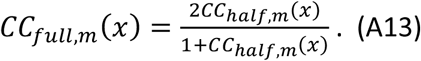

### Relationship between correlations

Let us assume that we have a map with the Fourier coefficients *F*_*o*_(*s*). The observation was made for the true map with the Fourier coefficients *F*_*t*_(*s*). And we have a model describing the true map with the Fourier coefficients *F*_*c*_(*s*). We assume that noise on the observations *F*_*n*_(*s*) is additive as well noise and signal are uncorrelated::

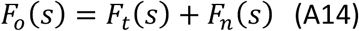

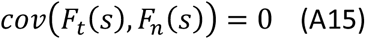

We also assume that noise in the observation is uncorrelated with the Fourier coefficients from atomic model (*Cov*(*F*_*c*,_ *F*_*n*_) *=* 0). Correlation between observed and calculated Fourier coefficients calculated within thin resolution shells is:

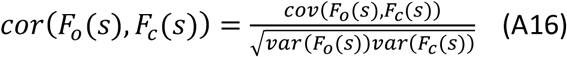

Since we assume that the correlation between observed noise and atomic model is zero we can write:

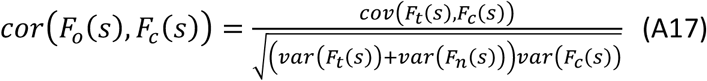

Correlation between observed and “true” Fourier coefficients can be written as:

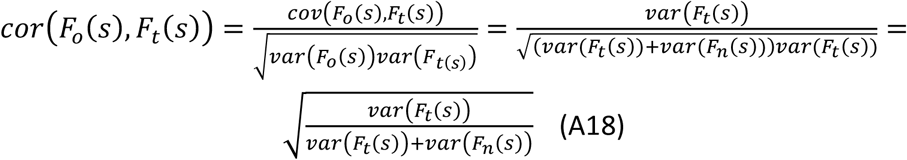

If we multiply the numerator and denominator of A17 by 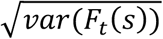 then we can write:

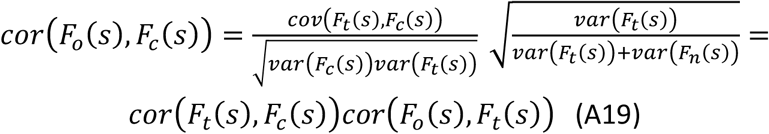

In practice we do not know “true” Fourier coefficients. However, if we can assume that we have two independent data sets (i.e. independent half data reconstructions) then we can use the expression (Rosenthal and Henderson, 2003)

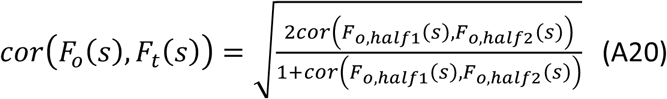

Therefore, we can write:

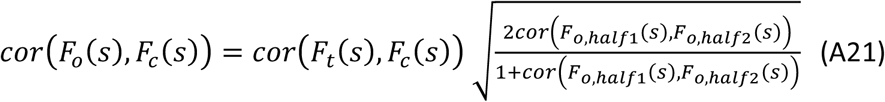

Although formulas are derived for correlations calculated in Fourier space, under above assumptions (uncorrelatedness of the noise and true and the noise and model) they are valid also for real space correlation.

## Appendix B

### Calculation of variances and covariances using the data

Let us assume that we have *N* observations and they are made for “true” maps. Noise is additive and uncorrelated with each other and with the signals. We also have half data reconstructed maps for each map. Thus:

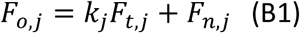

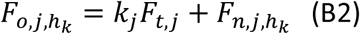

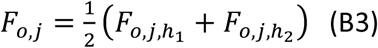

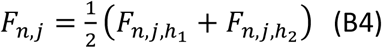

The noise components between half maps are uncorrelated, they have 0 mean and they have the same variance (i.e 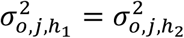).

Variances are calculated within resolution bins. This is described in a number of papers (Murshudov, 2016; Rosenthal and Henderson, 2003). Covariances between different maps within resolution bins are calculated using the formula:

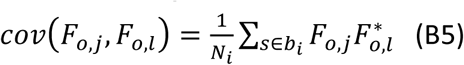

Where *N*_*i*_ is the number of Fourier coefficients within the resolution bin *b*_*i*_. Then the covariances are:

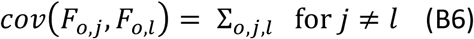

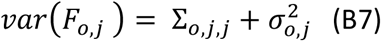

And using the half maps:

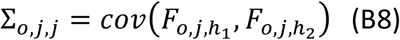

Using B7 and B8, variance of the noise for each map is calculated. It should be noted that when maps are being fitted into each other, the covariance matrix should be recalculated at every cycle. Also, the covariances should be adjusted to account for the effect of a mask (Chen et al., 2013).

## Appendix C

### Derivation of likelihood function and posterior probability distribution

Let us assume that we have *N* observed maps with Fourier coefficients - ***F***_***o***_(*s*) *=* (*F*_*o*,1_(*s*),*F*_*o*,2_(*s*),…., *F*_*o,N*_(*s*)). Each of *F*_*o,j*_ (*s*) is a complex number, i.e. it has two components – real and imaginary. Let us assume that these observations have been made for *N* “true” maps - ***F***_***t***_(*s*) *=* (*F*_*t*,1_(*s*),*F*_*t*,2_(*s*), …, *F*_*t,N*_(*s*)). In practice, the number of “true” maps could be less than the number of observed maps.

We assume that the underlying signals in those maps are related. For instance, those maps can be liganded-unliganded protein complexes, molecules in slightly different conformations, or related but not exactly the same macromolecules. Let us assume that for each “true” map we have a model – usually an atomic model from which we can calculate Fourier coefficients accounting for the nature of the experiment: ***F***_***c***_(*s*) *=* (*F*_*c*,1_(*s*),*F*_*c*,2_(*s*), …., *F*_*c,N*_(*s*)). We further assume that noise in the observations is additive, independent and with zero mean normal distribution.

We also assume that the conditional probability distributions of Fourier coefficients of the maps given the Fourier coefficients of true signals are Gaussian. Because of the central limit theorem this assumption holds in practice with sufficient accuracy.

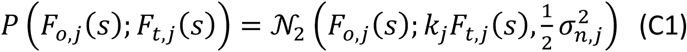

where 𝒩_2_ denotes two-dimensional normal distribution with mean equal to *k*_*j*_ (*s*)*F*_*t,j*_ (*s*) and variance equal to 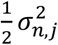. *k*_*j*_ is the scale parameter for the “true” map number *j* implying that “true” signal is blurred with a position independent point spread function before/during observations and/or data processing. Under an assumption that blurring is with an isotropic Gaussian kernel then *k*_*j*_ can be expressed in a form of Gaussian with a B value, 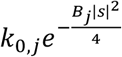. We will further assume that the “true” signals are on the same coordinate frame, however, observations may have been made for rotated and translated molecules. Then the probability distribution of individual Fourier coefficients will have the form:

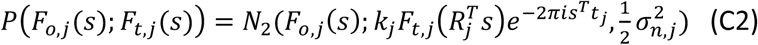

To get the total conditional probability distribution of observed Fourier coefficients all individual components are multiplied. Then, we can transfer transformations to the observed Fourier coefficients. To do this, it is assumed that variances of noise are the same on the surface of each sphere with a radius |*s*|. We also ignore correlation between different Fourier coefficients after transformation:

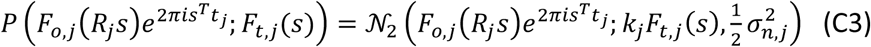

The probability distribution of the “true” Fourier coefficients given atomic model is also Gaussian, justification of which can be found in (Luzzati, 1952). Since “true” maps are related, we need to account for the relationship between different maps. We assume that the distribution of all “true” maps given all atomic models is Gaussian. This form of the distribution can be derived using the same technique used by Luzzati or the central limit theorem:

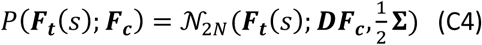

Where the subscript 2*N* signifies 2*N* dimensional normal distribution. ***D*** is a diagonal matrix formed by scale factors between calculated and “true” Fourier coefficients and **Σ** is the matrix of covariances

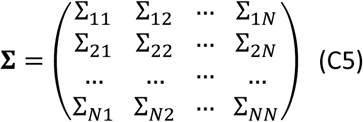

with Σ_*ij*_ =< (*F*_*t,i*_ − *D*_*i*_*F*_*c,i*_)(*F*_*t,j*_ − *D*_*j*_*F*_*c,j*_)* > to be estimated using data and atomic model. We also assume that observations are conditionally independent on model if the true map is known. In other words, if we know the true map, then atomic models would not say anything more about observations. Then the joint probability distribution of observed and “true” Fourier coefficients can be written as (Murshudov, 2016):

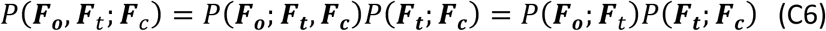

Since both distributions on the right-hand side are Gaussian, their product also will be Gaussian. In a multivariate Gaussian probability distribution, both marginal (integrating out some of the random variables) and conditional probability distributions of one subset given another subset of random variables are also Gaussian distributions (Eaton, 2007). To fully specify a Gaussian distribution, we need its mean vector and the covariance matrix.

*Likelihood function* is derived by integrating out the “true” unknown Fourier coefficients from the joint probability distribution of observations and “true” Fourier coefficients. I.e. it is a marginal probability distribution of observed Fourier coefficients. Since we know that the resultant probability distribution will be Gaussian with the mean and covariance matrix equal to the corresponding terms of the joint probability distribution of observed and “true” Fourier coefficients, we only need to find these terms. Since we know that:

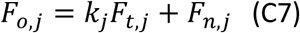

Therefore:

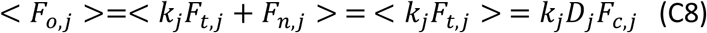

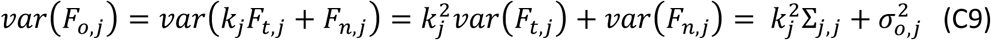

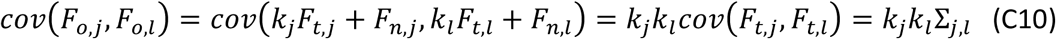

These fully specify the probability distribution of observed Fourier coefficients given calculated one. In practice, we cannot estimate all *k*_*j*_ without additional information. Relative values, *k*_*i*_*k*_*j*_^−1^, can be estimated using pairs of observed maps.

Coming back to our matrix/vector form using short notations ***F***_***o***_, ***F***_***t***_, ***F***_***c***_ for ***F***_***o***_(*s*),***F***_***t***_(*s*),***F***_***c***_(*s*) for clarity, we have

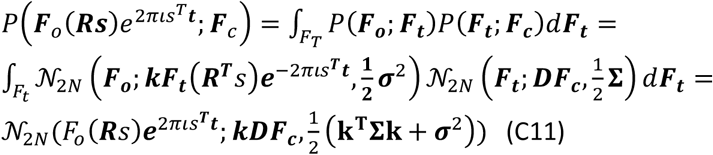

Where **Σ** is the covariance matrix of “true” map without blurring and **Σ**_*o,s*_ *=* ***k***^*T*^**Σ*k*** *=* ***k*Σ*k*** is the covariance matrix calculated using observed maps including half maps (Appendix B), ***k*** is a diagonal matrix formed by the scale factors of “true” Fourier coefficient. In the absence of models (***D*** → 0), the probability distribution will have the form:

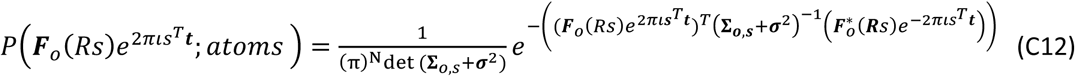

where *atoms* signifies that observations are made for a molecule that consists of atoms, but we do not know their positions.

The negative log likelihood function including all Fourier coefficients has the form:

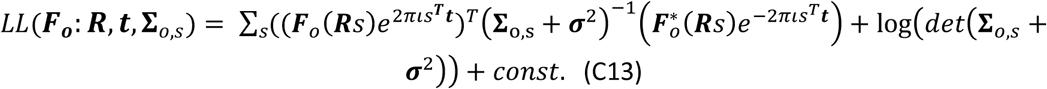

The formula C13 is used in EMDA to estimate rotations and translations of maps into each other as well as for magnification refinement. EMDA uses a special case of this, the two-observation case to fit two maps into each other. In general, ***R*** is a rotation matrix. However, if we relax this condition then we can also account for relative magnification of maps. If the only difference between maps is the relative isotropic magnification, then ***R*** will become a diagonal matrix where diagonal elements are relative magnification parameter.

*Posterior probability* of “true” Fourier coefficients given observations and atomic model is a conditional probability distribution of “true” maps given observations and model parameters:

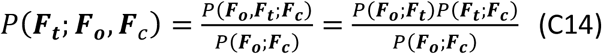

Again, the conditional probability distribution of a subset of random variables given another subset of variables in multivariate Gaussian distribution is also a Gaussian distribution (Eaton, 2007). So, we need to find the mean and covariance matrix. We know that the logarithm of a Gaussian distribution is a quadratic function. Argument that maximises this function is the mean of the random variable and the second derivative of this function with respect to the random variable we are interested in is related to the covariance matrix:

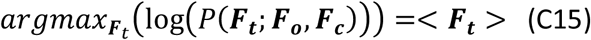

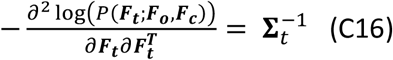

We need to find the argument that maximises the following function and its second derivative:

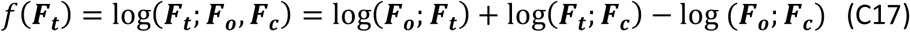

Since the third term on the RHS does not depend on ***F***_***t***_ it can be ignored. We can also ignore normalisation coefficients in the probability distributions, because they depend on covariances not on the “true” Fourier coefficients.

In the following treatment, we will use the fact that all involved matrices are symmetric. The covariance matrix is symmetric by its nature, and the rest of the matrices are diagonal and therefore symmetric.

So, we need to get the derivatives of (after ignoring terms independent on ***F***_*t*_):

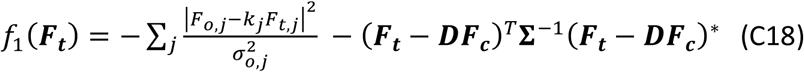

We can write:

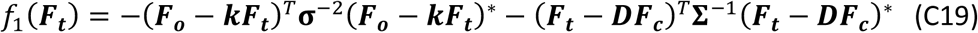

To find the maximum we need to solve:

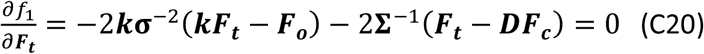

It can be conveniently solved:

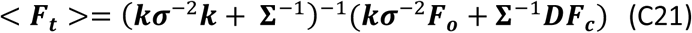

It shows that the mean value of the posterior probability distribution is a linear combination of observed and calculated Fourier coefficients with suitable weights.

Using the properties of matrices and their inverses we can write these formulas in a more convenient way:

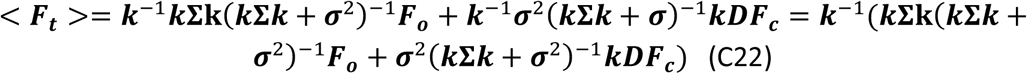

A special case of this when there is one model and one observation is considered in (Yamashita et al., 2021).

When there are no atomic models then ***D*** → *0* and the formula becomes:

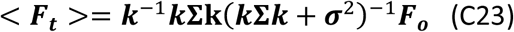

Further we denote **Σ**_***o***,*s*_ *=* ***k*Σ*k*** that can be estimated using the observed data:

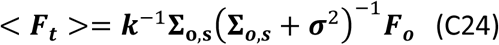

For completeness we also give the covariance matrix of the posterior probability distribution of the “true” Fourier coefficients (this can be used for estimation posterior noise variance and covariances in the calculated maps):

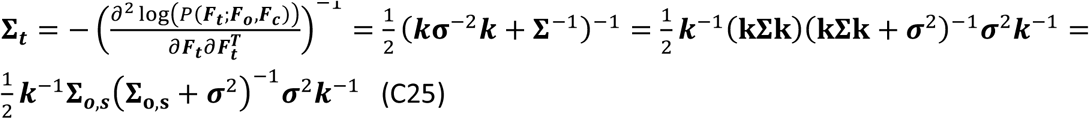

Since not all components of ***k*** can be estimated using the observations only, for current calculations we replace the elements of ***k*** with the standard deviations of the observed signal 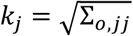 (explained in (Yamashita et al., 2021)). If ***k***_***1***_ is a diagonal matrix formed with 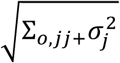 then we can write:

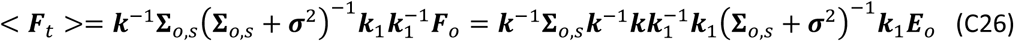

Here we used the notation 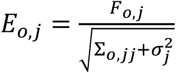. We recognise that ***ρ***_*s*_ *=* ***k***^−1^**Σ**_*o,s*_***k***^−1^ is the correlation matrix between “true” maps, 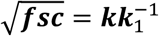 is the diagonal matrix formed with square root of fullmap FSCs, 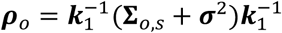 is the correlation matrix between observed maps. Now we can write:

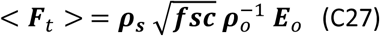

When there is only one map then this formula gives normalised and *fsc* weighted map. Next, we consider the case when N = 2. Then we can write the formulas in an explicit form:

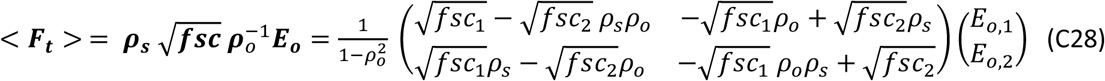

And

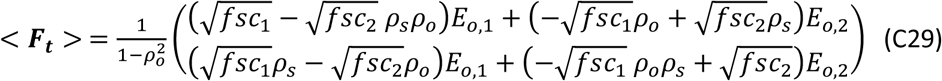

It must be stressed that since correlations are calculated using observed maps and when signal to noise ratio is very small then this estimation can vary dramatically. Therefore, for accurate estimations we may need to improve the estimation of the correlations, especially those between signal components, for example using smoothening or using prior knowledge derived from the PDB.

## Appendix D

### Use of normalized and weighted maps in local correlation calculation

In order to compare local correlations calculated using various maps, they need to be weighted appropriately in the same way.

The normalized expected map for a single map according to Bayesian interpretation is

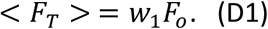

*F*_*o*_ is the observed Fourier coefficients, and 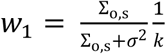 in which Σ_*o,s*_ and ***σ***^2^ are the covariance and the noise variance in the fullmap estimated using half maps in resolution bins as explained in Appendix B. *k* is a scale factor that associated with distortions of the true signal such as blurring. In the current implementation, *k* is replaced with the standard deviation of the observed signal (i.e. 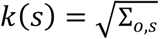) to yield

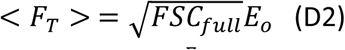

where 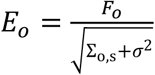.

While FSC-type weighting dampening down the noise, the normalisation works as a position independent deblurring operation.

Similar to D1, weights can be assigned on the calculated map as follows:

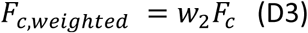

where *F*_*c*_ are calculated Fourier coefficients. The weights on calculated Fourier coefficients should be selected to dampen high resolution frequencies as in the weighted observed maps. Otherwise, the variance contribution of calculated high-resolution Fourier coefficients will reduce the correlation making it incomparable to that calculated for observed maps. We would also like to remove overall B value effect as in (D2). This way correlation in observed maps calculated using half maps will be comparable to that calculated between observed and calculated maps. To achieve this, we chose 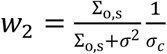 where *σ*_*c*_ is the standard deviation of calculated Fourier coefficients estimated in the same resolution bins as Σ_*o,s*_. Choosing such weights is equivalent to scaling *F*_*c*_ and *F*_*o*_ by making their variances equal, i.e.,

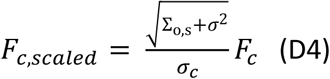

and using the calculated map with the following weights

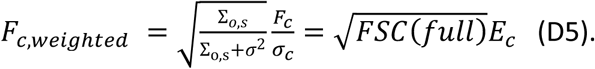

