## Supplementary materials for "EMDA: A Python package for Electron Microscopy Data Analysis"

### Relationship between scaling and blurring

Let us assume that there are two maps - $\psi_1(x)$ and $\psi_2(x)$ with corresponding Fourier coefficients - $F_1(s)$ and $F_2(s)$. Usually, to compare these maps or to calculate differences between them these maps are scaled, i.e. one of the Fourier coefficients is multiplied with resolution dependent scale factor, $k(s)$. I.e. $F_1(s)$ and $k(s)\ F_2(s)$ and/or their inverse transformations compared. According to the convolution theorem multiplying Fourier coefficients with a resolution dependent scale factor is equivalent to averaging the density at each point using the kernel derived from the inverse Fourier transformation of the scale factor:

$$\mathcal{F}^{-1}\big(k(s)F_2(s)\big) = \int_{y\in R^3} \psi_2(y)K(x-y)d^3y$$

Where $K(x) = \mathcal{F}^{-1}\big(k(s)\big)$. For this reason, often deblurring and scaling are used interchangeably. Thus, scaling in Fourier space is the same as blurring/sharpening in real space. If the scale factor is Gaussian then the blurring/sharpening kernel is also Gaussian. In practice, if there are more than one maps they can be scaled to each other.

### Kernel with different radii used in local correlation examples

Note that these are central slices of 3D kernels before normalisation.

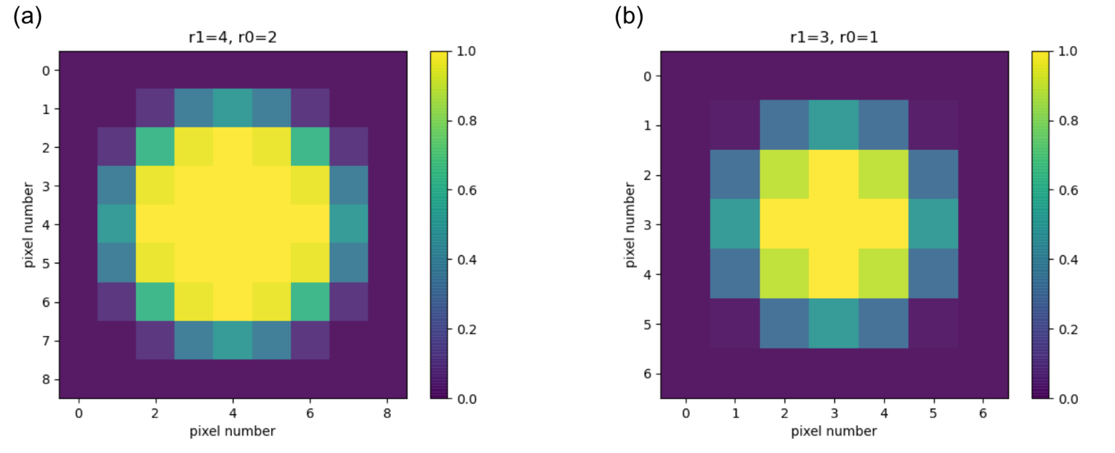

Fig. S1. 2-dimensional projections of 3D kernels used in correlation examples. a) kernel used in example 1, b) used in example 2.

### Effect of kernel size in local correlation

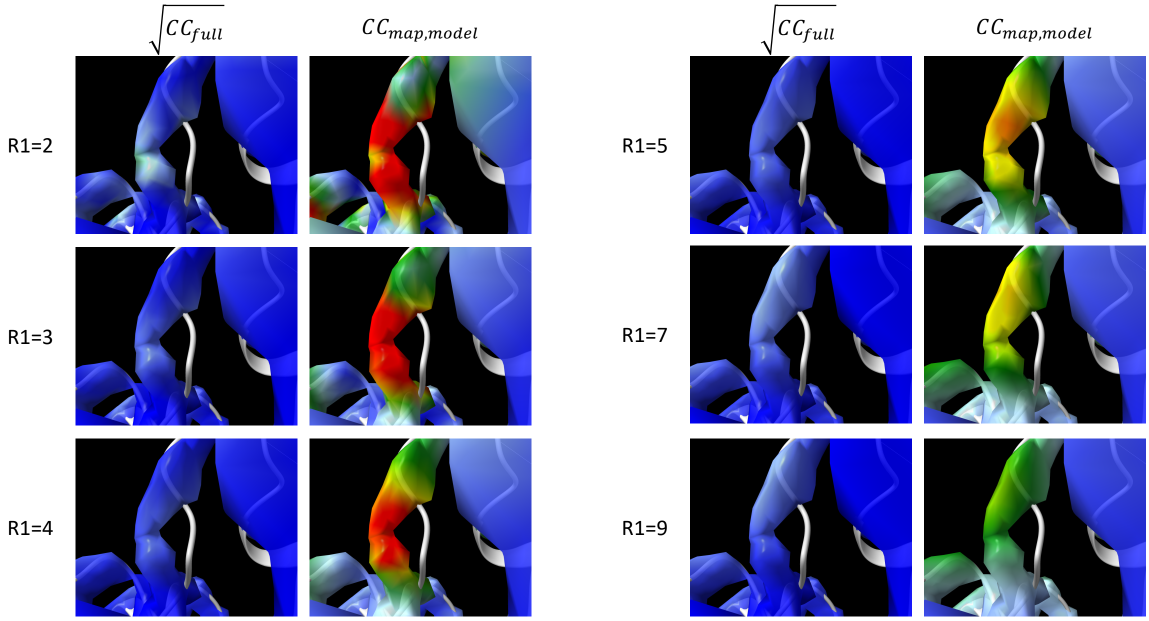


Local correlation example 1 in EMDA manuscript

Fig. S2. Effect of kernel size in local correlation example 1.

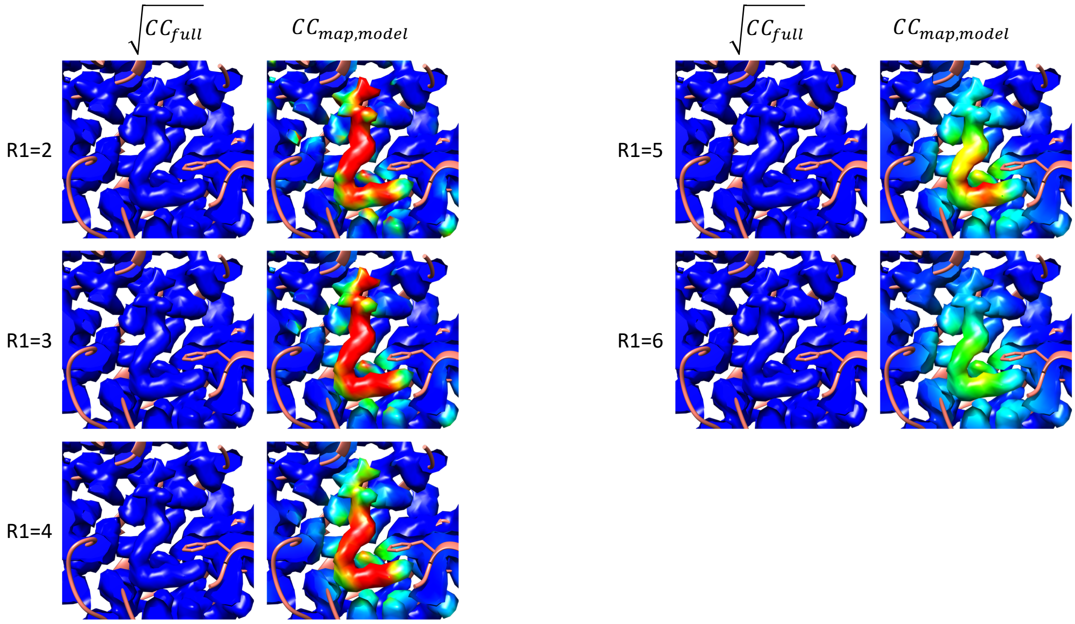


Local correlation example 2 in EMDA manuscript

Fig. S3. Effect of kernel size in local correlation example 2
